## Supplementary materials for "Evolution of sex differences in cooperation: the role of trade-offs with dispersal"

### Supplementary text

**1 - Quantifying the contributions of natal subordinates to cooperative provisioning**

To determine the rates at which helpers within their natal groups (i.e., all subordinate, and therefore non-breeding, group members still residing within the group in which they born) cooperatively provision the nestlings of the dominant pair, provisioning behaviour was recorded for all breeding attempts throughout the study period (2007 – 2016) in which the nestlings survived to the standard recording days. A video camera was placed on a low tripod below the nest to record the birds entering the nest on the mornings of the 6$^{th}$ to 12$^{th}$ days (inclusive) after the first egg in the focal clutch hatched (days that cover the period of highest nestling demand; nestlings fledge from the 17$^{th}$ day onwards). At least 5 days before video recording started, we caught every bird in the group apart from the dominant female (to avoid the risk of triggering abandonment) to confirm group composition and mark each bird's vent feathers with a uniquely positioned black dye mark to facilitate their identification on provisioning videos. On recording days, video cameras were set up at a standardized time following sunrise to record provisioning for approximately 3 hours per nest per day (yielding a total of 289 days of recording over the 177 monitored broods in groups with one or more helpers within their natal group; median of 2, range 1-6, days per brood). Within-nest cameras have confirmed that all nest visits in which the birds were not conspicuously carrying grass (events that were readily excluded from our provisioning rate calculations below), entail the visiting bird carrying a single food item to the nest and delivering it to the chicks [1]. The videos were subsequently transcribed to determine the timing of each provisioning visit, the duration of the visit (from the bird entering the nest to exiting), and the identity of the provisioning bird (using its unique vent mark and information from its leg rings and sex-specific bill colouration). Whenever the bill of the provisioning bird was visible on their approach to the nest, we also categorised the size of the food item being delivered; simply 'large' if it was evidently longer than the bird's bill or 'short' if it was not.

We used the resulting provisioning data set to calculate the number of provisioning visits that each group member made during each observation period (termed 'provisioning number' within our methods). In some cases, we were unable to reliably identify all group members, adding uncertainty to our provisioning number estimates. We therefore excluded any provisioning number estimates derived from observation periods in which there were five or more provisioning visits in which the feeder was of uncertain identity (72 provisioning number estimates excluded). To address the residual uncertainty in the provisioning number estimates derived from the observation periods containing one to four feeds by birds of uncertain identity, we set each bird's provisioning number estimate for these observation periods to be the integer midway between the lowest and highest possible provisioning number for that bird in that period given any partial identity information that was available for the feeders of uncertain identity. This yielded a final dataset containing 1,338 daily provisioning number measures for subordinates (helpers) of known sex within their natal groups for whom age was known with less than 180 days of uncertainty.

**2 - Additional Encounternet methods**

To calibrate the base stations, we attached two (transmitter) tags to a wooden pole with one of the tags' antenna positioned parallel to (vertical), and the other positioned perpendicular to (horizontal), the focal base station's antenna (to account for different antenna angles when the tags were on free-flying birds [2]. We then held the tags (on the pole) at a fixed distance of two meters from each base station and at a height of 1.70 meters for two minutes, resulting in 48 logged pulses with RSSI values for each base station. We then calculated the mean RSSI value logged by each base station ($mean_{Basestation}$) and the mean RSSI value over all base stations ($mean_{RSSI}$). For the 10 base stations for which $mean_{Basestation}$ was outside the range of $mean_{RSSI}$ ± standard deviation (SD), we adjusted the signal strength (RSSI) value of all logs from these base stations during analysis by adding $mean_{RSSI}$-$mean_{Basestation}$ (to avoid an over or underestimation of the distance between tagged birds and the focal base stations). We did not formally calibrate the individual tags (as well as the base stations), but we did verify qualitatively that all tags were yielding broadly comparable signal strengths at a fixed distance from a given base station prior to deployment. To ensure that the times at which tags were logged at different base stations could be compared over short timescales (to infer tag movements), we regularly checked that the clocks of all base stations were synchronised (yielding a typical deviation of between 0 and 1 seconds; maximum 6 seconds). Base station batteries were changed every 10 days, before they were discharged. Tag signal strength seemed to stay relatively constant during the period of data collection.

We discarded all logs of a given tag from (i) the day on which the tag was deployed and the following day [3] (to allow time for the tagged bird to get used to the tag and recover from any capture-related disturbance), and (ii) a short period following any attempt to catch a member of the tagged bird's home group (from the time of the capture attempt to the end of the following day; given the potential for capture attempts to impact the birds' movements). A given tag's logs were then processed in moving windows of 15 seconds and in time steps of 5 seconds. For each of these 15-second windows, we attempted to assign the tagged bird a 'best estimate' location following a conservative set of rules depicted as a flow diagram in Figure S3 and explained in the figure legend. Five birds were assigned to known locations (i.e., locations other than 'unknown' in Figure S3) for a low proportion of their total tracked time (<30%), most likely due to tag failure, and so were excluded from our analyses. The average proportion of time that the remaining 27 tagged birds were assigned known locations was 73%. Note that with the rule set for location assignment (Figure S3), a tag having weak transmission strength (e.g., due to tag error or signal obstruction) would not cause it to be falsely assigned 'non-home' locations (that might later be interpreted as prospecting), because assignment to 'non-home' locations required that the tag was logged with a stronger signal strength at a non-home group than the home group (Figure S3). Consecutive 15 second windows assigned to the same location were combined to represent 'time periods' for which our best estimate location was on a given territory ('unknown' locations [see Figure S3 legend] were removed from the tag's location time series prior to this step, as they could occur while the bird was at any location due to the bird's microenvironment impeding signal transmission).

*A priori* we set the distance threshold to detect extra-territorial forays to > 250m as this approach renders it highly likely that the focal bird itself is > 125m away from the centre of their home territory, because (given the workflow in Figure S3) its signal strength must have been stronger at a base station receiver > 250m away from home than it was at any other receiver in the array, including the one at the centre of its home territory. As the mean (± SE) distance between the centres of neighbouring territories is 93.7 m (± 4.56 m) in our study population, this minimum plausible distance of 125m from the centre of the tagged bird's home territory should ensure that 'forays' detected using this 250m distance threshold will typically have involved movements beyond the territory-centres of the tagged bird's neighbouring groups.

**3 - Impact of unknown dispersal events on age-specific dispersal probabilities of natal subordinates**

A small number of unringed adults born outside our study population do enter our study groups each year, suggesting that a small number of adults born within our study population probably also disperse beyond its boundaries each year [4]; unobserved dispersal events that this analysis will therefore miss. As males disperse significantly further than females in this population [4], and accordingly males are also more likely than females to enter our study population from outside [4], it would seem that males are more likely to be making undetected dispersals beyond the boundaries of our study site than females are. The key finding from this analysis, that males show a higher age-specific probabilities of dispersal than females is therefore if anything likely to underestimate the true magnitude of this sex differences in dispersal probability.

**4 - Sex difference in the strategy used to acquire dominance via dispersal**

As subordinate males and females are equally likely to inherit natal dominance (Main text Figure 2), the observed male-bias in age-specific dispersal probability (Main text Figure 1a) appears to reflect a sex difference in the *strategy* used to acquire dominance via dispersal, rather than in the *incidence* of acquiring dominance via dispersal. The data supports this: when subordinates dispersed from their natal groups, males were more likely than females to disperse into non-breeding subordinate positions in other groups (46 of 56 observed natal male dispersals; 19 of 32 observed natal female dispersals; $\chi^2 = 4.16$, $p = 0.041$, Table S10), where they might eventually win dominance or disperse again [4]. Whereas when females dispersed from their natal groups, they were more likely than dispersing males to disperse directly into a dominant position (10 of 56 dispersals for males; 13 of 32 dispersals for females; $\chi^2 = 4.16$, $p = 0.041$). Thus, males disperse more frequently than females, but employ a more complex dispersal strategy to acquire dominance often involving staging as a subordinate in other groups prior to dominance acquisition elsewhere, while females disperse less frequently but are more likely to do so directly into dominant positions elsewhere.

### Supplementary Figures

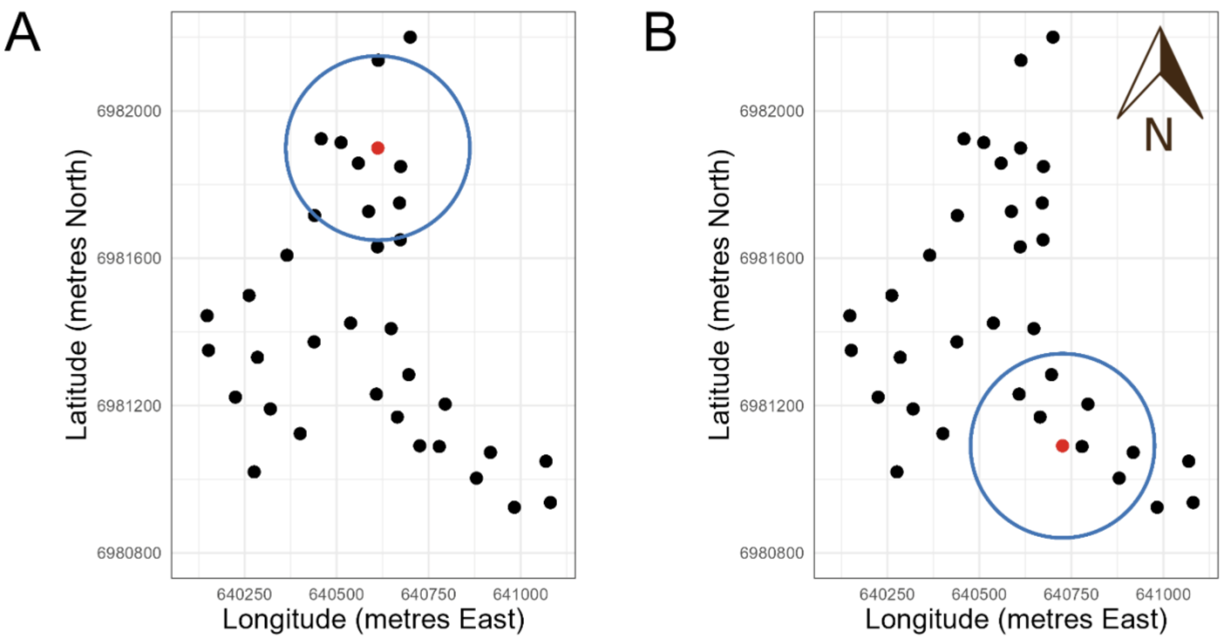


**Figure S1**. The Encounternet receiver "base station" array that was used for detecting forays. Panel **A** and **B** show the same simplified map of the study site with the locations of the 35 base stations (black and red dots), most of which were placed in the centre of distinct sparrow weaver territories (see methods). The x and y axes present longitude and latitude (respectively) in metres East and metres North of a given arbitrary location and thus also provide the scale for these maps. In each panel, a circle of 250 meters radius around a focal base station (red dot) is illustrated with a blue line; the only difference between the panels being the location of the focal base station. A single foray was defined as a continuous run (in time) of location estimates which suggested that the bird's closest base station was >250 m away from the centre of its home territory (i.e., outside blue circle) for at least 15 seconds. As the mean (± SE) distance between the centres of neighbouring territories was 93.7 m (± 4.56 m), the forays detected with this approach will typically have involved movements *beyond* the centres of the territories of neighbouring groups. This conservative approach will minimise the chance that a resident bird's territorial interactions with its neighbouring groups along their shared territory boundary are incorrectly interpreted as extra-territorial prospecting, but is likely to underestimate the true incidence of extra-territorial prospecting by excluding more local forays. The lack of base stations placed within the territories of study groups in the regions outside our core study population will also have left this approach underestimating true foray rate (and likely mean foray distance too). The land to the East and West of the presented array contains no other sparrow weaver territories within the pictured area (and so the focal birds will not have been conducting forays to groups living in those areas). However, there are a small number of widely spaced territories to the North of the array and several also lie close to the array to the South. Prospecting movements into these active territories will therefore not have been logged by our array. Spatial heterogeneity in the probability of true forays being detected by the array will have been accounted for in our statistical models of prospecting rate as all included social Group ID (i.e., territory ID) as a random term.

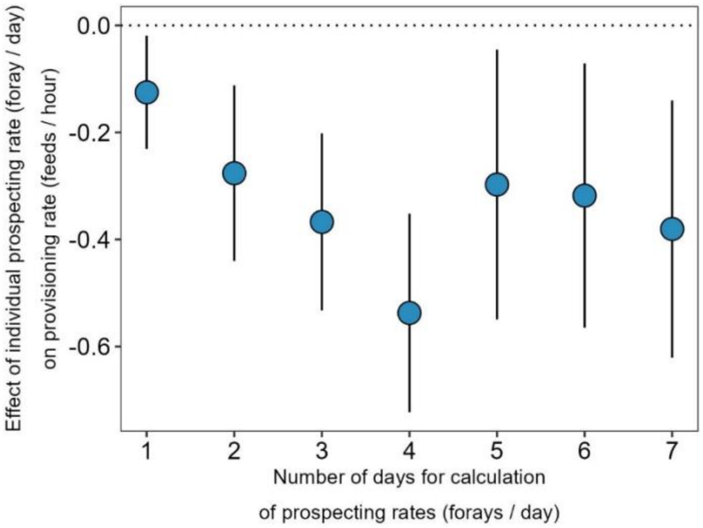


**Figure S2**. The effect of changing the number of days over which an individual's prospecting rate was calculated (x axis; prior to the day on which provisioning rate was measured) on the effect size estimate for the effect of an individual's prospecting rate (forays / day) on its cooperative provisioning rate (feeds / hour). Our analyses within the main paper calculated the prospecting rate over the 3 days prior to the measurement of provisioning rate, as we thought it plausible *a priori* that any energetic or stress-related costs of prospecting might accumulate and be evident over this timescale. However, on recognising that this decision is somewhat arbitrary we sought to verify, via this sensitivity analysis, that the detected negative covariance between the two traits wasn't particular to this choice of time window. The analysis confirms evidence of negative covariance between the two traits over a range of time windows. The initial steady increase in the effect size as the length of the time window considered increases could reflect (i) the timescale over which accumulated costs arising from recent prospecting impact cooperative behaviour, and/or (ii) that, given the modest rate at which prospecting forays occur, the shortest time windows may simply give a poorer-quality estimate of the focal bird's overall true rate of prospecting. Dots and error bars represent mean model estimates ± SE for the effect of prospecting rate on provisioning rate from the model presented in Table S9 when calculating each bird's prospecting rate over different time windows.

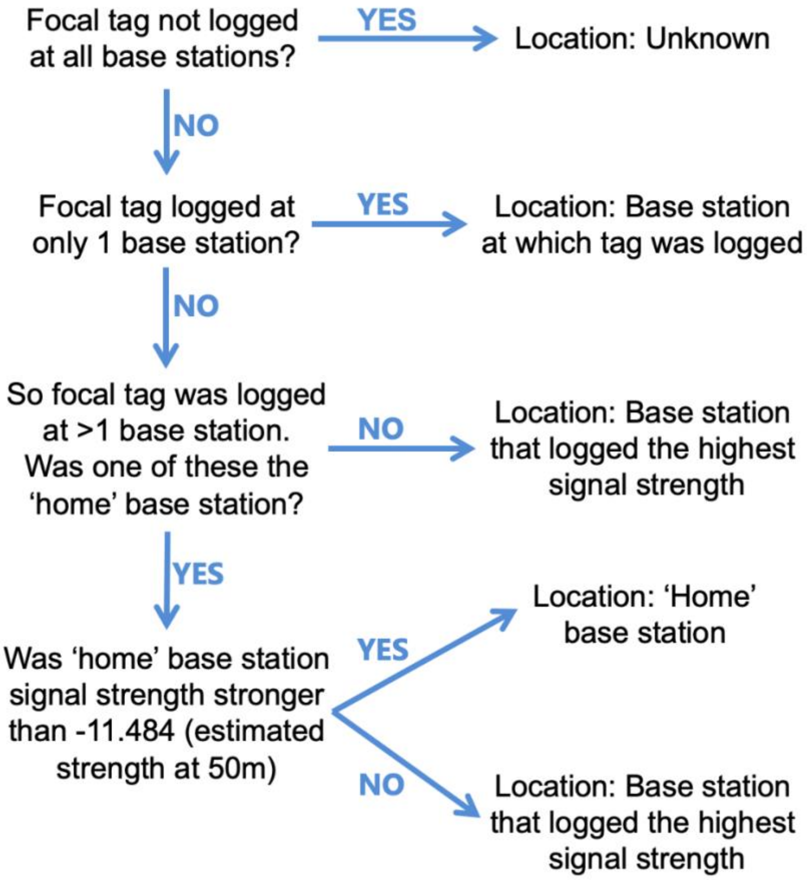


**Figure S3**. Decision tree applied to assign 'best estimate' locations to tagged birds using the logs from the Encounternet receiver base station array (see Methods). For each 15-second window in the time series of logs for a given bird, we attempted to assign the tagged bird a 'best estimate' location via the following set of rules depicted by the flow diagram here. First, if there were no logs at all for the focal tag during the focal time window, we noted the bird's location as 'unknown' (we did not draw spatial inferences from such 'unknown' location events as they could reflect the bird being in a microenvironment that impeded signal transmission, such as thick cover). Second, if there were logs for the focal tag from just one base station (indicating that the tag was out of reception range from all other base stations), we assigned the tagged bird the location of the base station at which it was logged. Third, if there were logs for the focal tag from more than one base station but none of them were the tagged bird's home base station, we assigned the tagged bird the location of the base station whose logs had the highest mean signal strength. Note that the highest mean signal strength could nevertheless have been weak in this case (e.g., if the bird was well beyond the boundary of our base station array, having prospected out of the study area, it's tag might be logged with only a weak signal strength at the base station closest to it on the array periphery; hence us terming these 'best estimate' locations). Fourth, if there were logs for the focal tag from more than one base station but one of them was the bird's home base station, (i) if the home base station signal strength was stronger than -11.484 (the estimated signal strength at 50m; see Figure S5) we assigned the tagged bird the home base station location, (ii) if this was not the case, we assigned the tagged bird the location of the base station with the strongest mean signal strength.

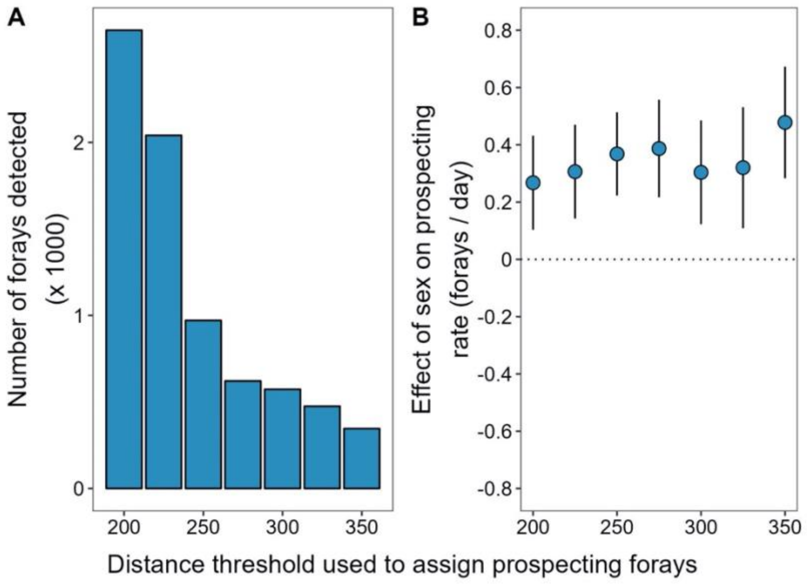


**Figure S4**. Sensitivity analysis to assess the effect of the 'distance threshold' set during the foray detection process (see Methods) on (**A**) the total number of detected 'forays' and (**B**) the effect size estimate for the sex difference in prospecting rate (forays / day; males relative to females). (**A**) A 'foray' was only considered to have occurred if the base station receiver that the focal bird was estimated to be closest to (i.e., its 'best estimate' location) was further than a set distance threshold away from the base station at the centre of the bird's home territory (see Methods). *A priori* we set this distance threshold to be 250 m as this approach renders it highly likely that the focal bird itself is > 125 m away from the centre of their home territory (see Methods in the main paper for the rationale). As the mean (± SE) distance between the centres of neighbouring territories is 93.7 m (± 4.56 m) in our study population, this minimum plausible distance of 125 m from the centre of the bird's home territory should ensure that 'forays' detected using a 250 m distance threshold will typically have involved movements beyond the territory-centres of the tagged bird's neighbouring groups. This approach should thereby minimise the chance that the bird's territorial interactions with its neighbours along their shared boundary while 'at home' are misclassified as extra-territorial forays. Reducing the distance threshold below 250 m will progressively increase the risk of such misclassifications; the likely cause of the marked increase in the number of 'forays' detected when threshold distances of 225 m and 200 m are used (panel **A**), while increasing the distance threshold above 250 m may yield an excessively conservative approach that substantially underestimates the incidence of 'true forays' by failing to capture those that occur over shorter distances. (**B**) The effect that changing this set distance threshold has on the estimated effect size (± SE) for the overall sex difference in prospecting rate (shown here as the effect of being male relative to female), when re-running the model presented in Table S6 using different distance thresholds during the foray classification process. The effect was consistently estimated to be positive across the range of distance thresholds tested (i.e., males having higher prospecting rates than females) and, as expected, the magnitude of the estimated sex difference effect size tended to increase as progressively higher distance thresholds were set. This is to be expected as lower distance thresholds will tend with greater frequency to misclassify home-territory movements (in which no sex difference is expected) as 'forays', thereby obscuring to a greater degree our estimate of the sex difference in the true prospecting rate. All of the prospecting analyses presented and referred to in the main text used the 250m distance threshold, as this makes most sense *a priori* from attention to the biology of the bird (see Methods).

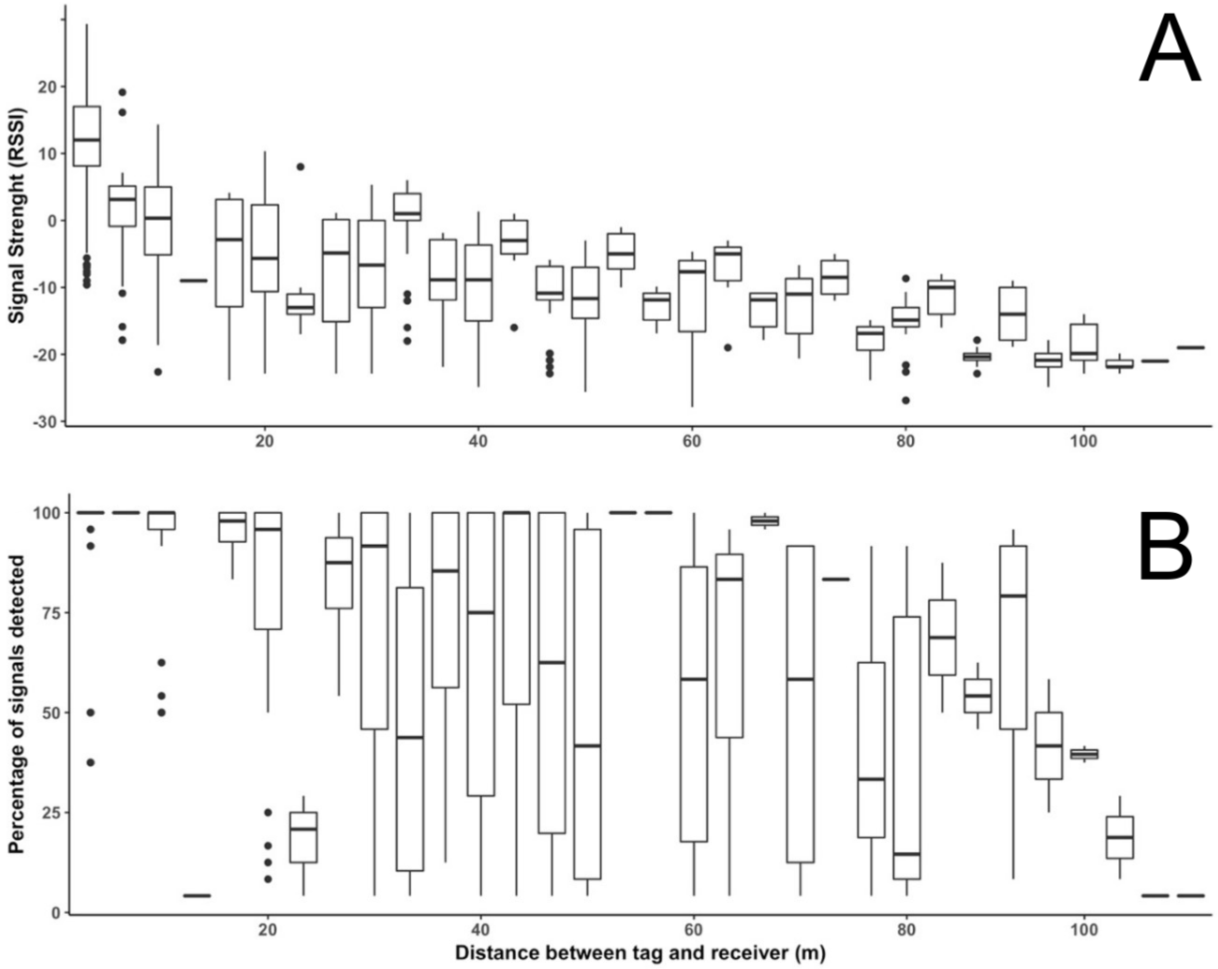


**Figure S5**. The Received Signal Strength Indicator (RSSI) values (**A**) and percentage of detected signals (**B**) both decreased with the distance between tags and base-station receivers in a field validation on our study site. Our workflow for using the distribution of signal strengths across our receiver array to allocate 'best estimate' locations for the tagged birds in each 15 second window, principally used information on the relative signal strength between receivers whenever tags were detected at multiple receivers simultaneously (see Figure S3 for details). However, to add an additional layer of conservatism to the assignment of 'non-home' locations in scenarios in which a tag was detected by the receiver in the centre of the tagged bird's 'home' territory as well as one or more receivers elsewhere, we also sought to estimate a threshold absolute signal strength that, if exceeded by the receiver on the home territory, would act as another indicator that the tagged bird was likely 'home'. To do this, we estimated the signal strength (RSSI)-distance relationship within our study site (panel A) using biologically realistic locations for the birds via the method described below, and then calculated the mean signal strength obtained at 50m distance from a receiver in this context for use as this threshold value (as the mean ± S.E. distance between neighbouring territory centres is just 93.7 m ± 4.56 m in our study population). This process yielded a threshold RSSI value of -11.484; so if a tagged bird was registered with an RSSI value above -11.484 at the receiver at the centre of its 'home' territory, the bird was conservatively assigned a 'home' location regardless of the signal strengths logged in other locations (see the final step in the Figure S3 workflow). We appreciate that the inherent variability within the RSSI-distance relationship (panel a; due to the effects for example of variation in tag height and signal obstruction via natural features on the study site) has two implications, and we do not consider either to be a problem. First, panel a highlights that tagged birds that are on their ‘home’ territory will not *always* be registered at their ‘home’ receiver at an RSSI >-11.484. In this scenario we expect the other conservative aspects of our workflows for (i) determining the ‘best estimate’ locations for birds (see Figure S3) and (ii) identifying forays from the properties of any ‘non-home’ locations (see main paper Methods) to ensure that the bird is not considered to be prospecting in this scenario. Second, panel A highlights that tagged birds could still be registered at their home receiver with an RSSI >-11.484 if they were 80m from that receiver (potentially further if at heights in excess of those tested in this field exercise; see below) and thus potentially just inside the territory of a neighbouring group. We consider the assignment of a ‘home’ location in this scenario appropriately conservative, as such locations could plausibly reflect routine territorial interactions between neighbouring groups rather than prospecting events. In order to characterise the relationship between RSSI and distance, we placed two tags on the end of a narrow wooden pole (one with its antenna oriented vertically, parallel to the base station antennae, and another with its antenna oriented horizontally, perpendicular to the base station antennae) and used the pole to move the tags among a series of locations on 10 transects, each starting from a different focal base station (itself hanging from a roost tree, just as the receivers in our array were). For each transect we assessed the signal strength of the tags first at 2m from the focal base station and then at locations at successive 10m intervals up to a maximum of 122m away from the focal base station. For one transect the first location was accidentally set at 8m from the focal base station and then sampled at 10m intervals up to 108m. At each location, we held the pole in position for four minutes, with tags located 1.70 meters above the ground for two minutes, and then on the ground for two minutes, to simulate tagged birds perching in vegetation and foraging on the ground; the two activities and approximate heights that dominate their time budgets.

### Supplementary Tables

**Table S1**. Coefficients and likelihood-ratio tests of Gaussian mixed model explaining variation in the provisioning visit duration of subordinates within their natal groups (response variable, originally in seconds, ln+1 transformed; n = 5,040 provisioning visits by 205 subordinates, 97 males and 109 females). The interaction between subordinate age and subordinate sex did not receive statistical support ($\chi^2_3 = 6.39$, p = 0.094) and was dropped from the full model to ease interpretation of single effect predictors. Residual variance = 0.262.

| Fixed effect | Estimate | SE[A] | 95% CI[A] | $\chi^2$ | df[A] | p |
|---|---|---|---|---|---|---|
| **Intercept** | 3.455 | 0.140 | 3.180, 3.729 | | | |
| **Subordinate sex** | | | | 16.46 | 1 | <0.001 |
| *Female* | — | — | — | | | |
| *Male* | -0.171 | 0.041 | -0.252, -0.090 | | | |
| **Subordinate age (years)** | | | | 8.08 | 3 | 0.044 |
| *< 1* | — | — | — | | | |
| *1-2* | -0.055 | 0.043 | -0.139, 0.028 | | | |
| *2-3* | -0.160 | 0.055 | -0.268, -0.052 | | | |
| *>4* | -0.049 | 0.101 | -0.247, 0.149 | | | |
| **Brood age** | | | | 32.75 | 6 | <0.001 |
| *6* | — | — | — | | | |
| *7* | -0.041 | 0.092 | -0.220, 0.139 | | | |
| *8* | -0.271 | 0.100 | -0.467, -0.076 | | | |
| *9* | -0.193 | 0.100 | -0.390, 0.003 | | | |
| *10* | -0.236 | 0.099 | -0.429, -0.042 | | | |
| *11* | -0.275 | 0.099 | -0.470, -0.081 | | | |
| *12* | -0.304 | 0.099 | -0.498, -0.110 | | | |
| **Brood size** | -0.095 | 0.050 | -0.193, 0.003 | 3.53 | 1 | 0.060 |

| Random effect variance | Estimate | # Levels |
|---|---|---|
| Individual ID | 0.486 | 205 |
| Social group ID | 0.000 | 31 |
| Breeding season | 0.005 | 8 |
| Clutch ID | 0.050 | 123 |

[A] SE = Standard Error, CI = Confidence Interval, df = degrees of freedom likelihood-ratio test.

**Table S2**. Coefficients and likelihood-ratio tests of binomial mixed model explaining variation in probability of provisioning a large food item by subordinates within their natal groups (n = 1,325 provisioning visits by 156 subordinates, 74 males and 83 females). The interaction between subordinate age and subordinate sex did not receive statistical support ($\chi^2_3$ = 0.31, p = 0.859) and was dropped from the full model to ease interpretation of single effect predictors. Model coefficients are shown in the link-function scale ('logit').

| Fixed effect | Estimate | SE[A] | 95% CI[A] | $\chi^2$ | df[A] | p |
|---|---|---|---|---|---|---|
| **Intercept** | 0.820 | 1.418 | -1.960, 3.599 | | | |
| **Subordinate sex** | | | | 1.43 | 1 | 0.231 |
| *Female* | — | — | — | | | |
| *Male* | -0.339 | 0.286 | -0.899, 0.222 | | | |
| **Subordinate age (years)** | | | | 3.57 | 3 | 0.312 |
| *< 1* | — | — | — | | | |
| *1-2* | 0.083 | 0.329 | -0.562, 0.728 | | | |
| *2-3* | 0.370 | 0.448 | -0.509, 1.249 | | | |
| *>4* | -15.044 | 114.487 | -239.434, 209.347 | | | |
| **Brood age** | | | | 7.73 | 6 | 0.258 |
| *6* | — | — | — | | | |
| *7* | -3.282 | 1.581 | -6.381, -0.184 | | | |
| *8* | -3.328 | 1.328 | -5.930, -0.725 | | | |
| *9* | -3.409 | 1.296 | -5.950, -0.868 | | | |
| *10* | -3.365 | 1.282 | -5.878, -0.853 | | | |
| *11* | -3.312 | 1.278 | -5.816, -0.807 | | | |
| *12* | -3.386 | 1.282 | -5.899, -0.872 | | | |
| **Brood size** | 0.119 | 0.338 | -0.543, 0.781 | 0.12 | 1 | 0.725 |

| Random effect variance | Estimate | # Levels |
|---|---|---|
| Individual ID | 0.391 | 156 |
| Social group ID | 0.396 | 27 |
| Breeding season | 0.000 | 7 |
| Clutch ID | 0.773 | 87 |

[A] SE = Standard Error, CI = Confidence Interval, df = degrees of freedom likelihood-ratio test.

**Table S3**. Microsatellite relatedness of subordinates and offspring (n = 487 relatedness measures for 205 subordinates, 111 males and 94 females). This table shows the result of a linear model explaining variation in microsatellite relatedness [5] between subordinates (males and females) and the offspring that they help to rear. In the model subordinate sex was included as a fixed effect predictor. For details of microsatellite genotyping, see Supplementary Text A in [6] and [7]. Residual variance = 0.041.

| **Fixed effect** | **Estimate** | **SE**[A] | **95% CI**[A] | **$\chi^2$** | **df**[A] | **p** |
|---|---|---|---|---|---|---|
| **Intercept** | 0.374 | 0.020 | 0.335, 0.413 | | | |
| **Subordinate sex** | | | | 0.16 | 1 | 0.691 |
| *Female* | — | — | — | | | |
| *Male* | -0.008 | 0.021 | -0.049, 0.032 | | | |
| **Random effect variance** | **Estimate** | **# Levels** | | | | |
| Clutch ID | 0.017 | 109 | | | | |

[A] SE = Standard Error, CI = Confidence Interval, df = degrees of freedom likelihood-ratio test.

**Table S4.** Coefficients and likelihood-ratio tests of binomial mixed model explaining variation in probability of subordinate dominance acquisition in the natal group when subordinates resided in the natal group at 1, 2, 3 and 4 years of age (n = 375 age-specific observations, 114 males and 105 females). The interaction between subordinate age and subordinate sex did not receive statistical support ($\chi^2_3 = 0.80$, $p = 0.850$) and was dropped from the full model to ease interpretation of single effect predictors. Model coefficients are shown in the link-function scale ('logit').

| Fixed effect | Estimate | SE[A] | 95% CI[A] | $\chi^2$ | df[A] | p |
|---|---|---|---|---|---|---|
| **Intercept** | -4.315 | 1.125 | -6.519, -2.111 | | | |
| **Subordinate age** | | | | 24.76 | 3 | < 0.001 |
| *1* | — | — | — | | | |
| *2* | 0.683 | 0.476 | -0.250, 1.616 | | | |
| *3* | 2.115 | 0.600 | 0.939, 3.292 | | | |
| *4* | 3.598 | 0.945 | 1.745, 5.451 | | | |
| **Sex** | | | | 0.29 | 1 | 0.592 |
| *Female* | — | — | — | | | |
| *Male* | -0.255 | 0.482 | -1.200, 0.690 | | | |

| Random effect variance | Estimate | # Levels |
|---|---|---|
| Social group ID | 3.953 | 35 |
| Breeding season of hatching | 0.398 | 6 |

[A] SE = Standard Error, CI = Confidence Interval, df = degrees of freedom likelihood-ratio test.

**Table S5.** Coefficients and likelihood-ratio tests of binomial mixed model explaining variation in probability of subordinate dominance acquisition in the natal group when subordinates resided in the natal group at 1, 2, 3 and 4 years of age, including events any in which dominance was acquired outside the natal group by founding a new group within territory previously held by the natal group (i.e., territorial budding [8]) (n = 375 age-specific observations, 114 males and 105 females). The interaction between subordinate age and subordinate sex did not receive statistical support ($\chi^2_3 = 0.03$, $p = 0.870$) and was dropped from the full model to ease interpretation of single effect predictors. Model coefficients are shown in the link-function scale ('logit').

| **Fixed effect** | **Estimate** | **SE**[A] | **95% CI**[A] | $\chi^2$ | **df**[A] | **p** |
|---|---|---|---|---|---|---|
| **Intercept** | -3.151 | 0.916 | -4.947, -1.355 | | | |
| **Subordinate age** | | | | 24.37 | 3 | < 0.001 |
| *1* | — | — | — | | | |
| *2* | 0.792 | 0.405 | -0.002, 1.587 | | | |
| *3* | 1.919 | 0.546 | 0.849, 2.988 | | | |
| *4* | 3.146 | 0.822 | 1.534, 4.758 | | | |
| **Sex** | | | | 1.89 | 1 | 0.169 |
| *Female* | — | — | — | | | |
| *Male* | -0.525 | 0.387 | -1.284, 0.234 | | | |

| **Random effect variance** | **Estimate** | **# Levels** |
|---|---|---|
| Social group ID | 2.033 | 35 |
| Breeding season of hatching | 2.557 | 5 |

SE = Standard Error, CI = Confidence Interval, df = degrees of freedom likelihood-ratio test.

**Table S6.** Coefficients and likelihood-ratio tests of Poisson mixed model explaining variation in prospecting rate (e.g., number of prospecting forays per day; n = 895 daily measures of prospecting rate from 27 tagged birds). No statistical support was found for the interaction between sex and provisioning ($\chi^2_1 = 0.97$, $p = 0.324$), which was removed from the final model. Model coefficients (Estimate) are shown along with standard errors (SE) and 95% confidence intervals (95% CI). Model coefficients are shown in the link-function scale ('log').

| **Fixed effect** | **Estimate** | **SE**[A] | **95% CI**[A] | $\chi^2$ | **df**[A] | **p** |
|---|---|---|---|---|---|---|
| **Intercept** | -0.546 | 0.199 | -0.936, -0.155 | | | |
| **Subordinate sex** | | | | 5.39 | 1 | 0.020 |
| *Female* | — | — | — | | | |
| *Male* | 0.368 | 0.145 | 0.083, 0.653 | | | |
| **Provisioning phase** | | | | 2.15 | 1 | 0.142 |
| *No* | — | — | — | | | |
| *Yes* | -0.144 | 0.094 | -0.328, 0.040 | | | |
| **Subordinate age** | 0.030 | 0.060 | -0.088, 0.148 | 0.23 | 1 | 0.631 |
| **Random effect variance** | **Estimate** | **# Levels** | | | | |
| Observation ID | 0.432 | 895 | | | | |
| Individual ID | 0.030 | 27 | | | | |
| Social group ID | 0.290 | 14 | | | | |

[A] SE = Standard Error, CI = Confidence Interval, df = degrees of freedom likelihood-ratio test.

**Table S7.** Coefficients and likelihood-ratio tests of Gaussian mixed model (with log-transformed response variable) explaining variation in duration of individual forays (log minutes; n = 971 prospecting forays from 27 tagged birds). No statistical support was found for the interaction between sex and provisioning ($\chi^2_1 = 1.14$, $p = 0.285$), which was removed from the final model. Model coefficients (Estimate) are shown along with standard errors (SE) and 95% confidence intervals (95% CI). Residual variance = 1.819.

| Fixed effect | Estimate | SE[A] | 95% CI[A] | $\chi^2$ | df[A] | p |
|---|---|---|---|---|---|---|
| **Intercept** | 2.405 | 0.288 | 1.840, 2.969 | | | |
| **Subordinate sex** | | | | 0.08 | 1 | 0.773 |
| *Female* | — | — | — | | | |
| *Male* | -0.060 | 0.235 | -0.521, 0.401 | | | |
| **Provisioning phase** | | | | 0.07 | 1 | 0.796 |
| *No* | — | — | — | | | |
| *Yes* | -0.029 | 0.111 | -0.247, 0.188 | | | |
| **Subordinate age** | -0.193 | 0.100 | -0.389, 0.002 | 3.95 | 1 | 0.047 |

| Random effect variance | Estimate | # Levels |
|---|---|---|
| Individual ID | 0.153 | 27 |
| Social group ID | 0.477 | 14 |

[A] SE = Standard Error, CI = Confidence Interval, df = degrees of freedom likelihood-ratio test.

**Table S8.** Coefficients and likelihood-ratio tests of Gaussian mixed model (with log-transformed response variable) explaining variation in distance of individual forays (log meters; n = 971 prospecting forays from 27 tagged birds). No statistical support was found for the interaction between sex and provisioning ($\chi^2_1 = 0.142$, $p = 0.707$), which was removed from the final model. Model coefficients (Estimate) are shown along with standard errors (SE) and 95% confidence intervals (95% CI). Residual variance = 0.059.

| Fixed effect | Estimate | SE[A] | 95% CI[A] | $\chi^2$ | df[A] | p |
|---|---|---|---|---|---|---|
| **Intercept** | 6.563 | 0.046 | 6.472, 6.654 | | | |
| **Subordinate sex** | | | | < 0.01 | 1 | 0.970 |
| *Female* | — | — | — | | | |
| *Male* | -0.002 | 0.040 | -0.080, 0.076 | | | |
| **Provisioning phase** | | | | 0.78 | 1 | 0.377 |
| *No* | — | — | — | | | |
| *Yes* | -0.018 | 0.020 | -0.057, 0.021 | | | |
| **Subordinate age** | -0.006 | 0.017 | -0.039, 0.027 | 0.14 | 1 | 0.708 |

| Random effect variance | Estimate | # Levels |
|---|---|---|
| Individual ID | 0.004 | 27 |
| Social group ID | 0.010 | 14 |

[A] SE = Standard Error, CI = Confidence Interval, df = degrees of freedom likelihood-ratio test.

**Table S9.** Poisson mixed model testing for a negative effect of prospecting rate (forays / day) on cooperative provisioning rate (feeds / hour). The data set comprised 34 daily measures of provisioning rate (each with a paired estimate of prospecting effort) from 13 subordinates (six males and seven females) in seven social groups. Prospecting rate was calculated in the three days leading up to the measurement of cooperative provisioning rate. Despite the modest sample size, the model recovered the significant negative main effect of prospecting rate on provisioning rate that is expected under the Dispersal trade-off hypothesis. The effects of subordinate sex, brood age and brood size were included as fixed effects only to ensure that uncontrolled independent effects of these variables were unlikely to be confounding the prospecting effect of interest. Given our use of a conservative full model approach throughout, these terms were retained in the model regardless of their significance. With regard to the interpretation of the effect of subordinate sex, see footnote B. Model coefficients (Estimate) are shown along with standard errors (SE) and 95% confidence intervals (95% CI).

| Fixed effect | Estimate | SE[A] | 95% CI[A] | $\chi^2$ | df[A] | p |
|---|---|---|---|---|---|---|
| **Intercept** | -2.321 | 1.356 | -4.979, 0.337 | | | |
| **Prospecting rate (forays/day)** | -0.367 | 0.165 | -0.691, -0.043 | 5.08 | 1 | 0.024 |
| **Brood size** | 0.415 | 0.357 | -0.286, 1.115 | 1.31 | 1 | 0.253 |
| **Brood age** | 0.201 | 0.099 | 0.008, 0.395 | 4.13 | 1 | 0.042 |
| **Subordinate sex** | | | | 1.93 | 1 | 0.165 |
| *Female* | — | — | — | | | |
| *Male* | -0.711 | 0.509 | -1.709, 0.287 | | | |

| Random effect variance | Estimate | # Levels |
|---|---|---|
| Individual ID | 0.471 | 13 |
| Social group ID | 0.000 | 7 |

[A] SE = Standard Error, CI = Confidence Interval, df = degrees of freedom likelihood-ratio test.
[B] While this lack of a significant effect of subordinate sex on cooperative provisioning rate (when simultaneously controlling for the effect of prospecting) also aligns with expectations under the Dispersal trade-off hypothesis, this result should be interpreted with caution as the power of this analysis to detect sex differences is likely low (data available for just six males and seven females).

**Table S10.** Coefficients and likelihood-ratio tests of binomial mixed model explaining variation in the probability that natal subordinate individuals emigrate to a subordinate position (n = 88 dispersal events; 46 of 56 observed natal male dispersals; 19 of 32 observed natal female dispersals). Model coefficients are shown in the link-function scale ('logit').

| Fixed effect | Estimate | SE[A] | 95% CI[A] | $\chi^2$ | df[A] | p |
|---|---|---|---|---|---|---|
| **Intercept** | 0.589 | 0.495 | -0.380, 1.559 | | | |
| **Sex** | | | | 4.16 | 1 | 0.041 |
| *Female* | — | — | — | | | |
| *Male* | 1.063 | 0.520 | 0.045, 2.081 | | | |

| Random effect variance | Estimate | # Levels |
|---|---|---|
| Social group ID | 0.000 | 30 |
| Breeding season of hatching | 0.266 | 7 |

[A] SE = Standard Error, CI = Confidence Interval, df = degrees of freedom likelihood-ratio test.

**References in Supplementary Materials**


1. Walker L. The evolution and regulation of cooperation in the wild. University of Exeter; 2015.

2. Rutz C, Morrissey MB, Burns ZT, Burt J, Otis B, Clair JJH. Calibrating animal-borne proximity loggers. Methods Ecol Evol. 2015;6: 656–667.

3. Clair JJH, Burns ZT, Bettaney EM, Morrissey MB, Otis B, Ryder TB. Experimental resource pulses influence social-network dynamics and the potential for information flow in tool-using crows. Nat Commun. 2015;6: 7197.

4. Harrison X, York JE, Young AJ. Population genetic structure and direct observations reveal sex-reversed patterns of dispersal in a cooperative bird. Mol Ecol. 2014;23: 5740–5755.

5. Queller DC, Goodnight KF. Estimating Relatedness Using Genetic Markers. Evolution. 1989;43: 258–275.

6. Capilla-Lasheras P, Harrison X, Wood EM, Wilson AJ, Young AJ. Altruistic bet hedging and the evolution of cooperation in a Kalahari bird. Sci Adv. 2021;7: 8980.

7. Harrison X, York JE, Cram DL, Hares MC, Young AJ. Complete reproductive skew within white-browed sparrow weaver groups despite outbreeding opportunities for subordinates of both sexes. Behav Ecol Sociobiol. 2013;67: 1915–1929.

8. Woolfenden GE, Fitzpatrick JW. The Inheritance of Territory in Group-Breeding Birds. Bioscience. 1978;28: 104–108.
